# Hyperimmune bovine colostrum containing lipopolysaccharide antibodies (Imm124-E) has a non-detrimental effect on gut microbial communities in unchallenged mice

**DOI:** 10.1101/2022.04.28.489964

**Authors:** Rachele Gore, Mitra Mohsenipour, Jennifer L Wood, Gayathri K Balasuriya, Elisa L Hill-Yardin, Ashley E Franks

**Author notes:** These authors have contributed equally to this work and share senior authorship (Elisa L Hill-Yardin, Ashley E Franks). **Correspondence:** Ashley Franks.

## Abstract

Enterotoxigenic *Escherichia coli* (ETEC) is a leading cause of bacterial diarrhea in travelers, military personnel and children in developing countries. Infection has the potential to cause long-term gastrointestinal dysfunction. Preventative treatments for ETEC-induced diarrhea exist, yet the effects of these treatments on gastrointestinal commensals in healthy individuals is unclear. Whether administration of a prophylactic preventative treatment for ETEC-induced diarrhea causes specific shifts in gut microbial populations in controlled environments is also unknown. Here we studied the effects of a hyperimmune bovine colostrum (IMM-124E) used in the manufacture of Travelan® (AUST L 106709) on gastrointestinal bacteria in healthy C57BL/6 mice. Using next generation sequencing, we aimed to test the onset and magnitude of potential changes to the mouse gut microbiome in response to the anti-diarrheagenic hyperimmune bovine colostrum product, rich in immunoglobulins against select ETEC strains (Travelan®, Immuron Ltd). We engineered changes in mouse fecal and cecal bacterial communities by delivering lipopolysaccharide (LPS) antibodies derived from bovine colostrum via dietary supplementation. Holstein Friesian and Jersey cows between 28- and 35-weeks’ gestation stimulated by subcutaneous delivery of three important pathogenic and antigenic determinants; LPS, flagella, and colonization factor antigen (CFA), produced a hyperimmune colostrum (IMM-124E) with demonstrated beneficial effects on health via modulation of metabolic pathways and immune function. We show that in mice administered colostrum containing LPS antibodies there was an increased abundance of potentially gut-beneficial bacteria, such as *Akkermansia* and *Desulfovibrio*, without disrupting the underlying ecology of the gastrointestinal tract. Compared to controls, there was no difference in overall weight gain, body or cecal weights or small intestine length following LPS antibody colostrum supplementation. Overall, dietary supplementation with colostrum containing LPS antibodies produced subtle alterations in gut bacterial composition of mice. Primarily, Travelan® LPS antibody treatment decreased the ratio of Firmicutes/Bacteroidetes in gut microbial populations in unchallenged healthy mice. Further studies are required to examine the effect of Travelan® LPS antibody treatment to engineer the microbiome in a diseased state and during recovery.

## 2 Introduction

Gut bacteria are part of a diverse population of commensal microorganisms that modulate host behavior and physiology. Endogenous bacteria influence the development of the gastrointestinal tract and immune system, the production of vitamins and release of neurotransmitters that regulate intestinal motility and fluid secretion. Microbes also assist the metabolism of otherwise indigestible dietary carbohydrates. Bacterial flora influence the structure and function of the gastrointestinal (GI) mucosal barrier and shape the broader physiology of the host, including impacts on the central nervous system and behavior via gut-brain axis pathways.

Enterotoxigenic *Escherichia coli* (ETEC) is a major cause of bacterial diarrhea in children in developing countries, military personnel and travelers (Jiang & DuPont, 2017; Kotloff et al., 2013; Shah et al., 2009; Walters et al., 2020). ETEC-induced diarrhea (also known as Traveler’s Diarrhea) is usually self-limiting, however, it can contribute to impairments in developmental processes or even mortality in young children (Troeger et al., 2018) and long-term post infection effects such as irritable bowel syndrome in travelers (Steffen, 2017). Alterations in gut microbiome profiles in response to infections with gram-negative pathogens responsible for Traveler’s Diarrhea have been identified (Walters et al., 2020; Youmans et al., 2015), yet the effects of ETEC antibodies on endogenous bacteria remain unclear.

Lipopolysaccharide (LPS) is a major component of the outer membrane of gram-negative bacteria such as *E. coli*. LPS is composed of a lipid A endotoxin connected via an oligosaccharide core to a hypervariable O-antigen polysaccharide (Raetz & Whitfield, 2002). The lipid A component is a potent activator of innate immune signalling via the Toll-like receptor 4/Myeloid differentiation factor 2 (TLR4/MD-2) complex (Matsuura, 2013). At high concentrations, lipid A induces acute inflammation resulting in fever, tachycardia, and septic shock in mammals, including humans (as reviewed in Alexander & Rietschel, 2001; Amorim et al., 2019). Orally ingested bovine immunoglobulins are an emergent therapeutic under investigation as both a preventative and a potential treatment of gastrointestinal dysfunction resulting from LPS-induced inflammation (Hutton et al., 2017; reviewed by Ulfman et al., 2018). Immunoglobulins targeting LPS can bind pathogens to prevent entry to host tissue via the mucosal epithelium. Furthermore, antibodies against LPS can modulate immune responses to enhance pathogen clearance and improve metabolic function by interacting with microorganisms in the gut (Adar et al., 2012; Mizrahi et al., 2012; Sears et al., 2017; Spalinger et al., 2018; reviewed by Ulfman et al., 2018). Although LPS influences host physiology via the microbiome (Lachmandas et al., 2016; Zhao et al., 2019), whether LPS antibody supplementation alters microbial populations is not known. Here, we investigated effects of dietary supplementation of colostrum containing LPS antibodies on gastrointestinal microbial richness and diversity in mice.

## 3 Materials and Methods

### 3.1 Preparation of colostrum and *E. coli* antibodies

Holstein Friesian and Jersey dairy cows at commercial dairy farms were immunized with either monovalent or polyvalent ETEC vaccines containing 0.5 mg of purified proteins administered subcutaneously in three 1 ml doses. The vaccines contain three important pathogenic and antigenic determinants; lipopolysaccharide (LPS), flagella, and colonization factor antigen (CFA), which collectively play roles in bacterial membrane stability, immune evasion, motility, and adherence. Three ETEC doses were administered prior to calving at 9-12 weeks, 6 weeks, and 1 month before calving. The monovalent vaccine used to immunize each animal contained the common ETEC strain; serotype O78, and the polyvalent vaccine contained a combination of serotypes (O6, O8, O15, O25, O27, O63, O114, O115, O128, O148, O153, O159) that were selected to cover the majority of *E. coli* that cause Traveler’s Diarrhea (Table 1). To produce a rapid and durable immune response, Montanide ISA 206 veterinary adjuvant (approved by the Australian Pesticides and Veterinary Medicines Authority) was used in preparation of the vaccine. Inoculation with these outer antigens activates a generalized immune response in the host animal to produce antibodies (mainly IgG) which recognize and bind with the bacterial cell-surface epitopes presented. These polyclonal antibodies can also cross-react with other similar gram-negative enteric pathogenic bacterial cell surface antigens. The Immuron Ltd hyperimmune bovine colostrum (IMM-124E) is a pasteurized, fat and lactose reduced spray dried powder which contains polyclonal antibodies targeting the endotoxin LPS and other bacterial components. IMM-124E contains 80% proteins, out of which approximately 35% to 40% are immunoglobulins. IMM-124E is harvested from the colostrum of dairy cows that have been immunized against the outer antigens, mostly LPS, of the most common strains of ETEC. In Australia, Travelan® is a listed medicine in the Australian Register for Therapeutic Goods (AUST L 106709) and is specifically indicated to reduce the risk of Travelers’ Diarrhea and reduce the symptoms of minor gastro-intestinal disorders.

**Table 1.** Enterotoxigenic *E. coli* (ETEC) strains in inactivated ETEC vaccines (Immuron Ltd.)

| SEROTYPE | STRAIN NO. | SOURCE | DATE OF ISOLATION |
| --- | --- | --- | --- |
| ETEC O6: H16 | B2C | USA | Pre-1971 |
| ETEC O8: H19 | C55 3/3c3 | USA | mid-1980's |
| ETEC O15: H4 | PE 595 | IMVS. Adelaide | Aust. Source |
| ETEC O25: H42 | E11881A | USA | 1986 |
| ETEC O27: HR | C1067-77 | USA | 1985 |
| ETEC O64: H- | PE 673 | IMVS. Adelaide | Aust. Source |
| **ETEC O78: H11** | H10407 | USA | Pre- 1973 |
| **ETEC O114: H21** | E20738/0 | UK | 1980 |
| **ETEC O115: H-** | PE 724 | IMVS. Adelaide | Aust. Source |
| **ETEC O128: H21** | EI 37-2 | USA | 1985 |
| **ETEC 148: H28** | B7A | USA | Pre- 1971 |
| **ETEC O153: H12** | E8772/0 | UK | Pre- 1980 |
| **ETEC O159: H-** | PE 768 | IMVS. Adelaide | Aust. source |

The hyperimmune bovine colostrum containing the Travelan® LPS antibodies listed in Table 1 used in this study are referred to as ‘Travelan® LPS antibody treatment’.

### 3.2 Mice used in the study

To assess for changes in mouse GI microbial populations following ingestion of the Travelan® LPS antibody treatment/placebo, twenty C57BL/6 male mice aged 4 weeks were imported from the Animal Resource Centre in Western Australia to RMIT University, Bundoora and randomly allocated to a placebo or treatment group.

### 3.3 Feeding protocol

Four-week-old mice were habituated for 2 days and randomly allocated to a placebo or treatment group. Mice were weighed prior to commencing the feeding protocol (days 0,1, 2) and at the termination of the feeding protocol (day 8). Researchers were not blinded to treatment.

Mice were not food-deprived prior to the feeding experiment. Standard chow (Specialty Feeds Irradiated Rat and Mouse diet cubes, Specialty Feeds, Glen Forrest, Western Australia) and water were available *ad libitum* during the 2 days prior to the implementation of 5 consecutive days of Travelan® LPS antibody treatment or placebo. In the group-housed cages, the standard chow was removed from 10 am to 3 pm for the 5 days of the feeding protocol. During the food restriction period, mice were placed in individual open-topped cages for 1 h twice a day and were given access to the LPS antibody treatment or placebo solutions. Mice within the placebo group were given access to 2 ml of freshly made 10% weight/volume Promilk solution, and mice in the treatment group were given access to 2 ml of freshly made 10% weight/volume Travelan® LPS antibody solution. The water-soluble powders were reconstituted before the start of each feeding session by dissolving 3 g of powder in 30 ml dH_2_0 and stirred slowly for up to 5 min on a magnetic stirrer to avoid damaging the proteins. A 30 mm diameter plastic petri dish containing either placebo or Travelan® LPS antibody treatment was attached to the floor of the cage with double sided tape to allow mice free access to the solution. After 1 hour, mice were returned to group housing cages, the remaining treatment/placebo solution was weighed, and the amount consumed by each mouse per day was calculated.

### 3.4 Sample collection

Fresh fecal samples were collected using sterile forceps, placed in individual sterile 1.5 ml Eppendorf tubes, snap frozen in liquid nitrogen and stored at −80 °C. Fecal samples were collected at the end of each isolation period twice daily for 2 days prior to commencing the feeding protocol, and then twice daily for the 5-day feeding protocol. Individual open-topped cages were cleaned using 70 % ethanol at the end of each isolation period.

At the end of the protocol, all 20 mice were culled by cervical dislocation (Ethics approval #1810, RMIT University) and final body weight recorded. Gut tissue was dissected, and anatomical measurements (cecal weight, length of small intestine and colon) were recorded. Cecal microbial content was transferred directly into sterile tubes, colonic content and colonic mucosal samples were collected aseptically from each individual mouse and snap frozen in liquid nitrogen for microbial analysis.

### 3.5 Microbial DNA extraction

Microbial genomic DNA was extracted from approximately 0.25 g of fecal, cecal and colonic samples using the QIAGEN DNeasy PowerSoil™ DNA isolation kit (QIAGEN, Hilden, Germany) as per manufacturer’s instructions. DNA concentrations were measured using Qubit™ dsDNA HS Assay kit and Qubit™ 3.0 Fluorometer (ThermoFisher Scientific; Invitrogen, MA, USA). All samples were stored at −30°C until required.

### 3.6 Quantitative PCR

Quantitative PCR (qPCR) was used to quantify total bacterial DNA copy number as an indicator of abundance. The primer pair 1114f-1275r, which targets the bacterial 16S rRNA gene (Denman & McSweeney, 2006) was used to detect bacterial communities. The qPCRs were run on a CFX Connect™ Real-Time PCR Detection System (Bio-Rad). Each 5-μl reaction solution contained 0.25 pM of each forward and reverse primer, 2.5 μL of SensiFAST™ SYBR® & Fluorescein mix (Bioline), sterile DNA-free water and 5 ng of sample DNA. Thermocycling conditions were 20 s at 95 °C, followed by 40 cycles of 95 °C for 3 s, and 61.5 °C for 30 s (Klevenhusen et al., 2011).

Reactions were followed by a melting curve increasing 1 °C every 30 s, from 60 °C to 99 °C. Bacterial copy number was quantified by using 1114f-1275r primers to amplify the 16s rRNA gene from *Escherichia coli* (DH5α). Standard curves were generated using triplicate 10-fold dilutions of the *E. coli* purified amplicon.

### 3.7 16S rRNA metabarcoding and bioinformatics

The V3-V4 hypervariable regions of the bacterial 16S rRNA gene (forward primer 5′ TCGTCGGCAGCGTCAGATGTGTATAAGAGACAGCCTACGGGNGGCWGCAG 3′; reverse primer 5′ GTCTCGTGGGCTCGGAGATGTGTATAAGAGACAGGACTACHVGGGTATCTAATCC 3’) was amplified on an Illumina MiSeq platform (2 × 300 bp). Raw, demultiplexed, FASTQ files were re-barcoded, joined, and quality filtered following the UPARSE operational taxonomic unit clustering pipeline (Edgar, 2016b) (USEARCH; http://drive5.com/uparse/). Joined paired-end reads were quality filtered by discarding reads with total expected errors >1.0. Reads that could not be assembled were discarded. Sequences were denoised and the -unoise3 command was used to generate exact sequence variants (ESVs). Taxonomic assignments were performed using the USEARCH SINTAX algorithm (Edgar, 2016a). Reference databases were created using the RDP_trainset_15 datasets available from UTAX (http://drive5.com/usearch/manual/utax_downloads.html). The minimum percentage identity required for an ESV to consider a database match a hit was 90%. Only ranks with a high enough confidence were retained. A phylogenetic tree was generated using MUSCLE (Edgar, 2004). ESVs identified as chloroplasts and mitochondrial DNA were removed from the data.

### 3.8 Statistical analysis

#### 3.8.1 Anatomical analysis

Statistical analyses were performed using GraphPad Prism v9.0.1 for Windows (GraphPad Software, San Diego, California USA). Cecal weight, small intestinal length and colon length taken at the time of culling were adjusted for individual body weight. Anatomical measurements were analyzed using two-tailed, Student’s *t*-test to assess for differences between Travelan® LPS antibody-treated and placebo-treated mice, and p< 0.05 was taken to indicate statistical significance. Data are presented in a box and whisker plot, displaying the mean, median, interquartile range and range of data.

#### 3.8.2. Microbial analysis

Statistical analyses were performed in the R statistical computing software environment (version 4.0.5) (R Development Core Team, 2014). We conducted alpha-diversity analysis to assess for differences in microbial communities across treatment groups. To determine whether there were differences in community structure between the placebo and Travelan® LPS antibody treatment microbial communities, we analyzed beta-diversity. Alpha- and beta-diversity analyses were performed on OTU-matrices rarefied to a depth of 10,000 reads using the ‘phyloseq’ (McMurdie & Holmes, 2013) and ‘vegan’ (Oksanen et al., 2013) packages.

Normality and variance homogeneity of the data were tested using the ‘shapiro.test’ and ‘bartlett.test’ functions. Alpha diversity metrics were assessed using t-tests or, where appropriate analysis of variance (ANOVA) testing with Tukey honest significant difference (HSD) test used for post-hoc testing. Non-metric multidimensional scaling (NMDS) ordinations were used to visualize community beta diversity. Dissimilarity matrices were generated using the UniFrac metric (Lozupone & Knight, 2005). Significance testing of beta diversity was carried out using permutational MANOVAs (PERMANOVA) tests of factors affecting community structure (Anderson, 2001).

Differential abundance testing of OTUs was performed using the DESeq2 extension available within the ‘phyloseq’ package (Love et al., 2014; McMurdie & Holmes, 2013). The differential abundance tests were performed on un-normalized community data as recommended by McMurdie and Holmes (2014), for each taxonomic rank to identify which individual ESVs were changing in response to Travelan® LPS antibody treatment. P values were corrected for multiple testing, using the Benjamini-Hochberg false discovery rate controlling procedure (Benjamini & Hochberg, 1995).

## 4 Results

To characterize the effects of colostrum containing LPS antibodies on the microbiome of unchallenged animals, mice were fed a placebo (milk powder) solution or a solution containing Travelan® LPS antibody treatment, and fecal microbes were collected to enable microbial population comparisons. Specifically, we aimed to test the onset and magnitude of the effect of hyperimmune bovine colostrum containing ETEC LPS antibodies on the mouse gut microbiome by assessing microbial richness and diversity in fecal, colon mucosal and cecal samples using deep sequencing methods.

### 4.1 Effects of LPS antibody-containing colostrum on gastrointestinal anatomy

Prior to assessing for changes in mouse GI microbial populations following LPS antibody colostrum supplementation, we first sought to determine if the treated mice exhibited any gross anatomical differences compared with placebo-fed mice. At the end of the feeding protocol, all 20 mice were culled, and anatomical measures (cecal weight, small intestinal length and colon length) were taken at the time of culling. These measures were then adjusted for individual body weight.

To determine if mice fed Travelan® LPS antibody treatment have altered gut structure compared to placebo-fed mice, body weight (Supplementary Figure 1) and gross anatomical measures of the gastrointestinal tract were recorded from freshly culled mice. All 20 mice consumed the milk or treatment solution each day of the experiment. No significant difference in the consumption of placebo/ Travelan® LPS antibody solution between treatment groups for the duration of the protocol was observed (Supplementary Figure 1). All mice in both placebo and LPS antibody colostrum supplemented groups gained weight during and after the feeding protocol (Supplementary Figures 2 and 3). There were no significant differences in cecal weight normalized to body weight between placebo-fed mice and Travelan® LPS antibody-treated mice (Supplementary Table 1; p=0.140). No significant differences were identified in small intestine length normalized to body weight between placebo-fed mice and Travelan® LPS antibody-treated mice (Supplementary Table 1; p=0.452). A trend towards increased colon length in LPS antibody-treated mice compared to placebo-fed mice (Supplementary Table 1, p=0.056) (Figure 1) was observed.

**Figure 1.**
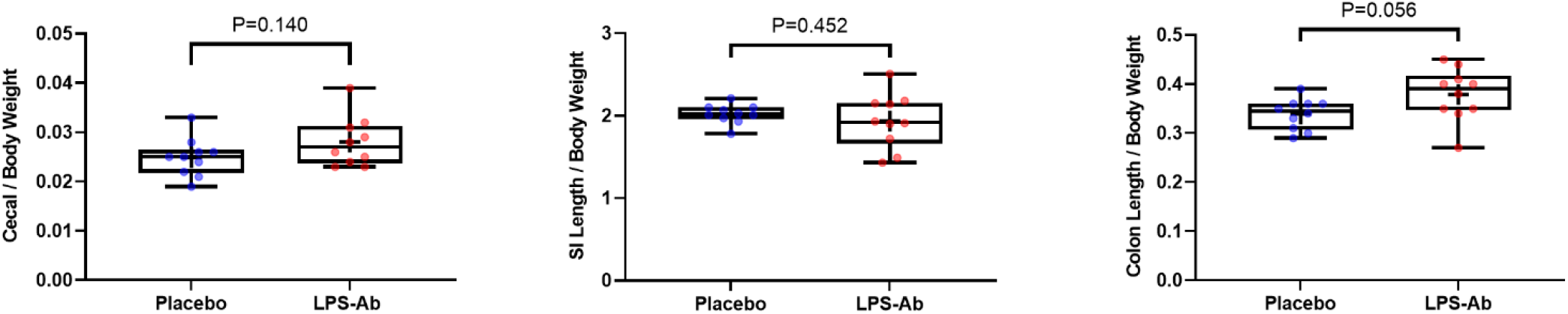
Gastrointestinal tract anatomical measures normalized to body weight. Left, mouse cecal weight (p=0.140); center, small intestinal (SI) length (p=0.452); right, colon length (p=0.056); as a proportion of total body weight. Placebo: Promilk 85, Tatura Milk Industries Ltd, Australia; LPS-Ab: Travelan® LPS antibody colostrum solution. Boxplots represent median, interquartile range and range of the data; cross inside the boxplot represents the mean.

### 4.2 Total bacterial quantification

To investigate the impact of Travelan® LPS antibody treatment on bacterial numbers, bacterial diversity and species richness, microbial samples from different regions of the gastrointestinal tract were collected from mice following completion of the feeding protocol.

We used quantitative PCR (qPCR) and next generation sequencing to investigate if Travelan® LPS antibody treatment induced changes to the diversity and species richness of bacterial communities from fecal, colonic mucosal and cecal samples. Our qPCR data showed that Travelan® LPS antibody treatment had no effect on bacterial cell numbers in any of the gastrointestinal community samples.

Although we initially saw an increase in bacterial cell number in fecal samples for both groups of mice, this number had begun to return to the initial values by the end of the feeding protocol, resulting in no significant difference in total copy number between the treatment groups, or when compared to before the feeding protocol commenced (Figure 2).

**Figure 2.**
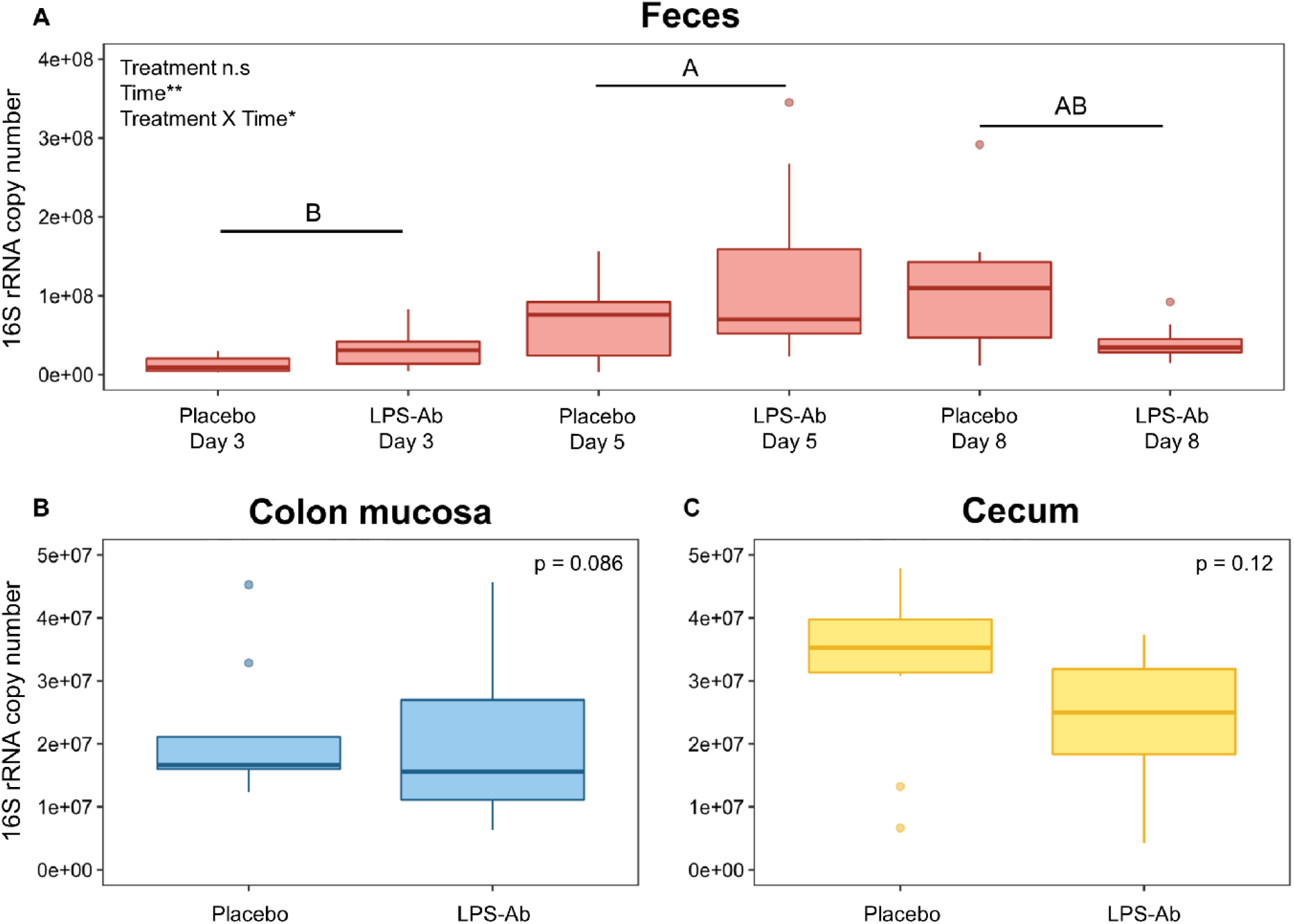
qPCR estimates of total bacterial cell number based on 16S copy number for feces, colon mucosa and cecum samples. Fecal samples (panel A) from days 3 (pre-treatment), 5 (early treatment) and 8 (final treatment) of the experiment. Letters indicate statistical difference in copy number between sampling times (Days). Bacterial numbers collected from days with different letters are significantly different (treatment effect P> 0.05, time effect (Collection Day) P < 0.05, two-way ANOVA, panel A). The total number of bacterial cells in mucosal (panel B) and cecal samples (panel C) at day 8 (end of experiment) were similar. Placebo: Promilk 85, Tatura Milk Industries Ltd, Australia); LPS-Ab: Travelan® LPS antibody treatment. Boxplots represent median and interquartile range; whiskers represent 1.5 x interquartile range (Tukey method).

### 4.3 Community richness and diversity

Across fecal communities and colonic mucosal content collected post-mortem, there was no significant effect of time or Travelan® LPS antibody treatment on bacterial diversity (Shannon-Wiener diversity), OTU (Operational Taxonomic Units) richness or evenness (Figure 3). However, community evenness was reduced in the cecal communities of mice supplemented with the Travelan® LPS antibody treatment when compared to controls (p=0.009, Figure 3).

**Figure 3.**
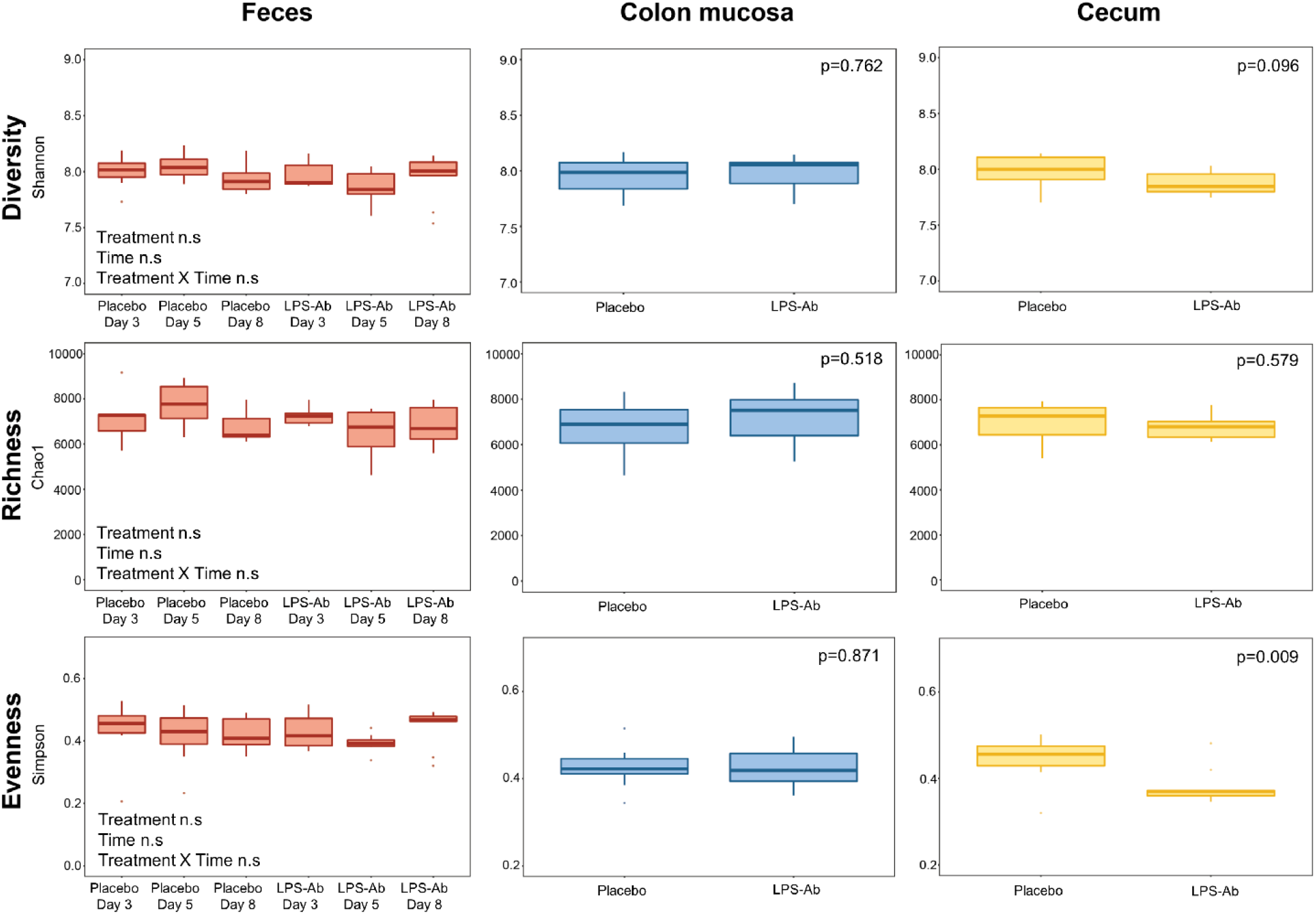
Bacterial community diversity metrics; Shannon diversity (top), Chao1 richness (middle) and Simpson’s evenness (bottom). Fecal samples collected pre-treatment (Day 3) and after treatment (Day 5 and 8). Day 3 represents the first day of feeding the mice with the Travelan® LPS antibody treatment (LPS-Ab)/placebo solution (n=10 placebo, n=10 LPS-Ab). Colon samples were collected post-mortem (Day 8) and include primarily mucosal content, with some fecal content n=10 placebo, n=10 LPS-Ab. Cecal samples were collected post-mortem (Day 8) n=10 placebo, n=10 LPS-Ab. Placebo: Promilk 85, Tatura Milk Industries Ltd, Australia); LPS Ab: Travelan® LPS antibody treatment. Boxplots represent median and interquartile range; whiskers represent 1.5 x interquartile range (Tukey method).

Travelan® LPS antibody treatment significantly impacted the structure of fecal communities as detected by Two-way PERMANOVAs (pseudo-*F* = 1.9313, p=0.0008). LPS antibody colostrum treatment also resulted in low-level but significant grouping of bacterial communities for both colonic mucosal (pseudo-*F* = 1.8485, p=0.0069) and cecal (pseudo-*F* = 1.3428, p=0.007) communities (Figure 4). For cecal communities, the strength of grouping due to the antibody treatment remained significant when data was presence-absence transformed (i.e., converted to 0s and 1s), suggesting that Travelan® LPS antibody treatment altered both bacterial abundance in addition to the species present in cecal communities (Figure 4).

**Figure 4.**
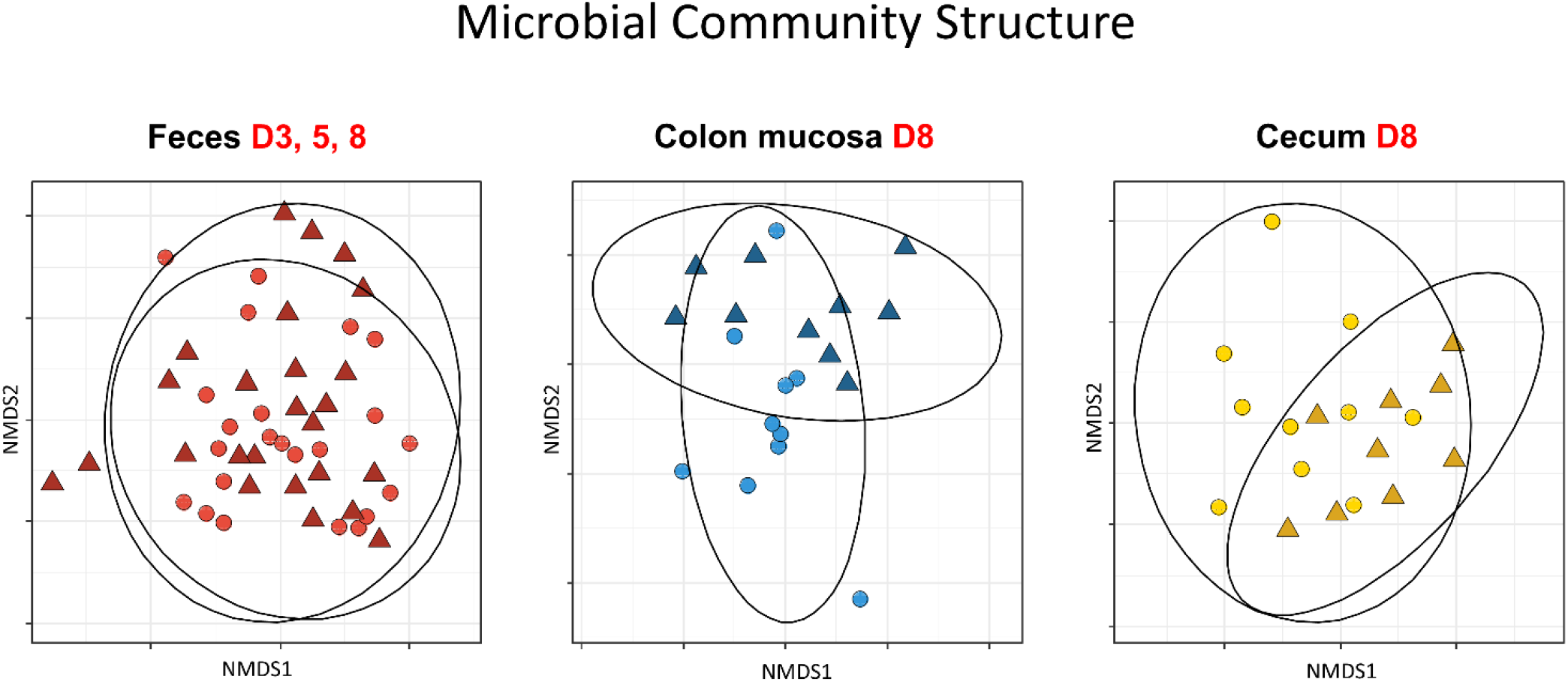
Non-metric multidimensional scaling (NMDS) ordination showing relationship between bacterial community structure from fecal samples from Days (D) 3, 5 and 8 of the experiment (left, pseudo-*F* = 1.9313, p=0.0008) and colonic mucosal samples (center, pseudo-*F* = 1.8485, p=0.0069) and cecum samples (right, pseudo-*F* = 1.3428, p=0.00709) from D8 (end of experiment). Circles = placebo; Triangles = Travelan® LPS antibody treatment.

In summary, these data suggest that Travelan® LPS antibody treatment had minimal to no impact on bacterial cell number, bacterial species richness or species abundance distribution (evenness) in fecal and colon mucosal samples. In contrast, in cecal samples there was a reduction in the relative abundance of the different species found in the Travelan® LPS antibody-treated samples compared to placebo-treated samples. Cecal samples from LPS antibody-treated mice exhibited a trend for an overall reduction in species diversity compared to controls. At the community level, small but significant shifts in community structure were observed for fecal, mucosal and cecal samples following treatment.

### 4.4 Travelan® LPS antibody treatment results in differentially abundant bacterial OTUs

Given that Travelan® LPS antibody treatment impacted bacterial community structure in fecal, mucosal and cecal samples, differential abundance testing of OTUs was performed for each taxonomic rank to identify which individual OTUs were changing in response to the treatment. Across fecal samples, the genera *Akkermansia* and *Enterorhabdus* were enriched in Travelan® LPS antibody treated communities and microbes within the genus *Clostridium* (Clade III) were reduced (Figure 5, Table 2). *Akkermansia* displayed the largest significant change in abundance of all genera irrespective of sample type. In cecal content communities, Travelan® LPS antibody treatment decreased the abundance of 8 bacterial genera, in particular; *Porphyromonas*, a producer of short-chain fatty acids (Guilloux et al., 2020), and *Parabacteroides* which contributes to bile acid metabolism and plays a role in reinforcing the enteric epithelium (Hiippala et al., 2020; Li et al., 2020) (Figure 7, Table 4). For mucosal communities, no individual OTUs or genera differed significantly in their abundance, however, there was a trend towards an increase in *Olsenella* and *Desulfovibrio*, and a decrease in *Clostridium* (Clade XVIII) (Figure 6, Table 3). At higher taxonomic levels (Family and Order) the Desulfovibrionales were significantly enriched Travelan® LPS antibody treated mucosal communities.

**Table 2.** Genus-level OTUs with altered abundances for fecal communities (NB: low abundance of *Enterorhabdus*; blue, a large change in abundance of *Akkermansia*; bold).

| RELATIVE<br>ABUNDANCE | LOG2FC | EFFECT<br>OF<br>LPS AB | ADJ<br>P-VALUE | PHYLUM | ORDER | GENUS |
| --- | --- | --- | --- | --- | --- | --- |
| **0.27** | **6.09** | Increase | **7.86E-06** | Verrucomicrobia | Verrucomicrobiales | ***Akkermansia*** |
| 0.07 | 1.03 | Increase | 7.44E-03 | Actinobacteria | Coriobacteriales | *Enterorhabdus* |
| 0.77 | -1.25 | Decrease | 2.56E-02 | Firmicutes | Clostridiales | *Clostridium_III* |

**Table 3.** No significant alteration to genus-level OTUs in colon mucosal communities.

| RELATIVE<br>ABUNDANCE | LOG2FC | EFFECT<br>OF<br>LPS AB | ADJ<br>P-VALUE | PHYLUM | ORDER | GENUS |
| --- | --- | --- | --- | --- | --- | --- |
| 0.20 | -3.73 | Decrease | 0.057 | Firmicutes | Erysipelotrichales | *Clostridium XVIII* |
| 0.21 | 4.23 | Increase | 0.057 | Actinobacteria | Coriobacteriales | *Olsenella* |
| 0.46 | 1.62 | Increase | 0.075 | Proteobacteria | Desulfovibrionales | *Desulfovibrio* |

**Table 4.** Altered genus-level OTUs for cecal communities (highest relative abundance in bold).

| RELATIVE<br>ABUNDANCE | LOG2FC | EFFECT<br>OF<br>LPS AB | ADJ<br>P-VALUE | PHYLUM | ORDER | GENUS |
| --- | --- | --- | --- | --- | --- | --- |
| **1.10** | **-1.64** | **Decrease** | **0.001** | **Bacteroidetes** | **Bacteroidales** | ***Porphyromonas*** |
| 0.26 | -2.34 | Decrease | 0.004 | Bacteroidetes | Bacteroidales | *Saccharicrinis* |
| 0.52 | -1.93 | Decrease | 0.005 | Bacteroidetes | Sphingobacteriales | *Parapedobacter* |
| 0.03 | -2.43 | Decrease | 0.010 | Bacteroidetes | Sphingobacteriales | *Pedobacter* |
| 0.35 | -3.47 | Decrease | 0.014 | Firmicutes | Erysipelotrichales | *Allobaculum* |
| 0.03 | -2.54 | Decrease | 0.014 | Bacteroidetes | Flavobacteriales | *Robiginitalea* |
| **3.55** | **-1.25** | **Decrease** | **0.014** | **Bacteroidetes** | **Bacteroidales** | ***Parabacteroides*** |
| 0.02 | -2.53 | Decrease | 0.040 | Bacteroidetes | Sphingobacteriales | *Sphingobacterium* |
| 17.55 | 1.12 | Increase | 0.061 | Firmicutes | Clostridiales | *Clostridium XIVa* |

**Figure 5.**
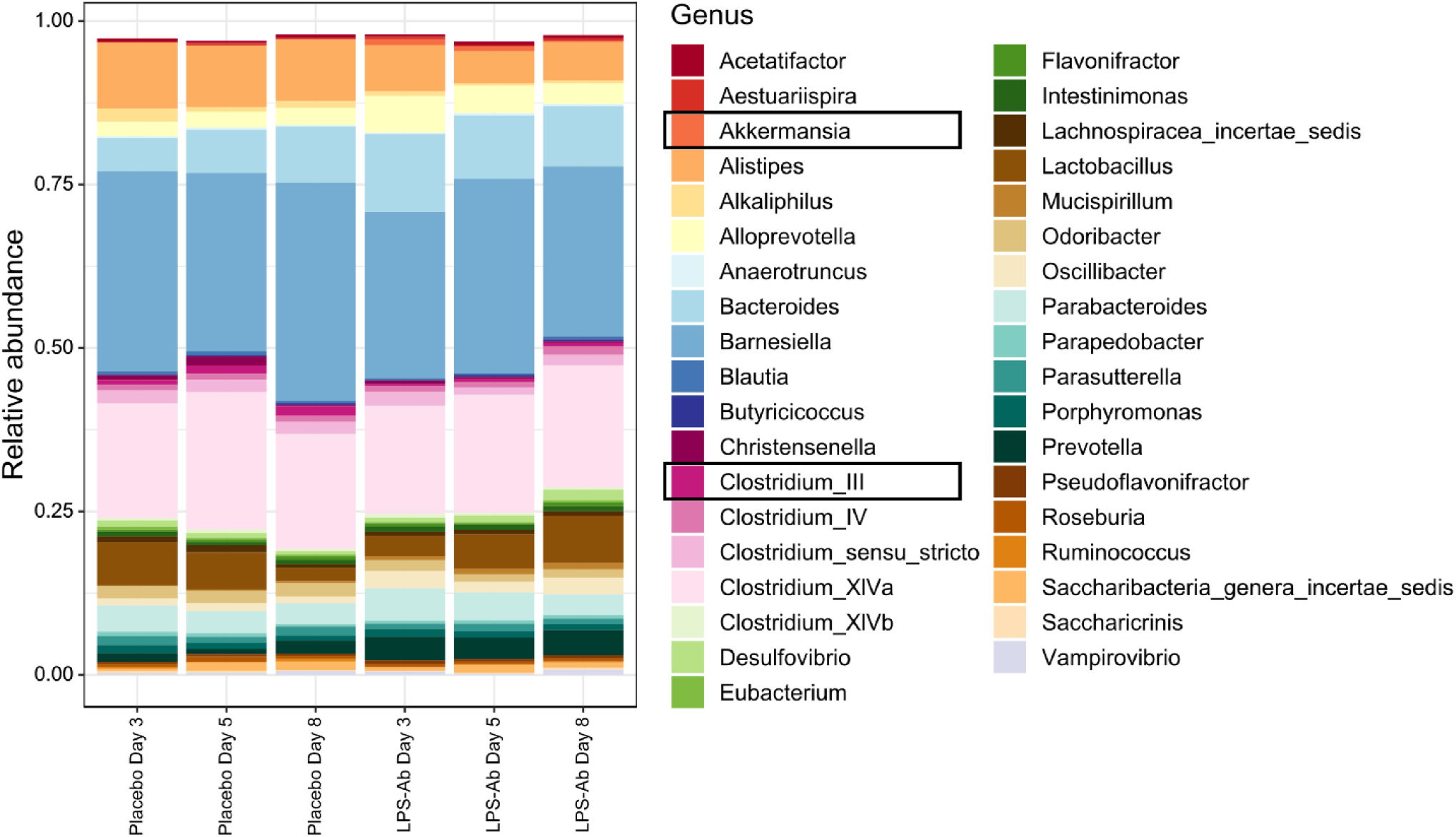
Descriptive bar charts of fecal communities from Travelan® LPS antibody-treated and placebo-treated communities. OTUs colored by genus. Y axis represents % abundance. 78 bacterial genera each with a mean abundance < 0.005% have been excluded for clarity. (Increased abundance of *Akkermansia* and decreased abundance of *Clostridium III* visualized here) (NB: due to the low level of abundance, *Enterorhabdus* is not depicted in the bar chart).

**Figure 6.**
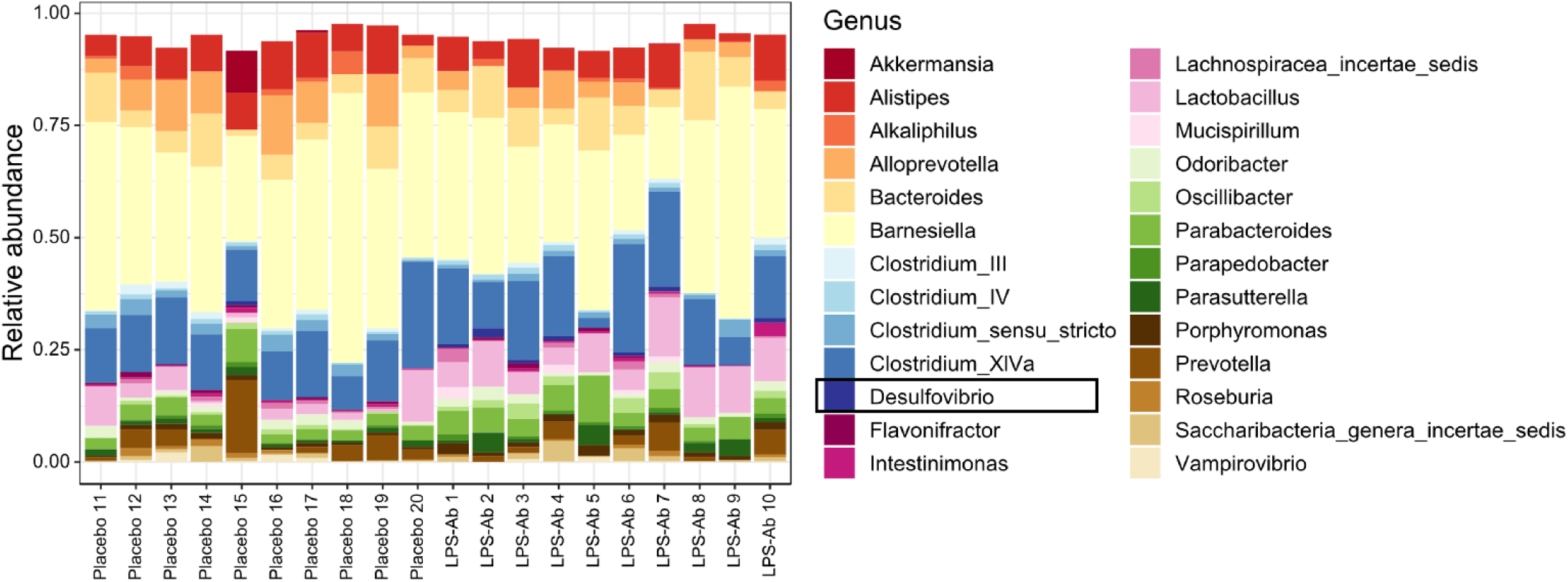
Descriptive bar charts of colon mucosal communities from Travelan® LPS antibody-treated and placebo-treated communities. OTUs have been colored by genus. Y axis represents % abundance. 76 bacterial genera each with a mean abundance < 0.005% have been excluded for clarity. (NB: due to the low level of abundance, *Clostridium XVIII* and *Olsenella* are not depicted in the bar chart).

**Figure 7.**
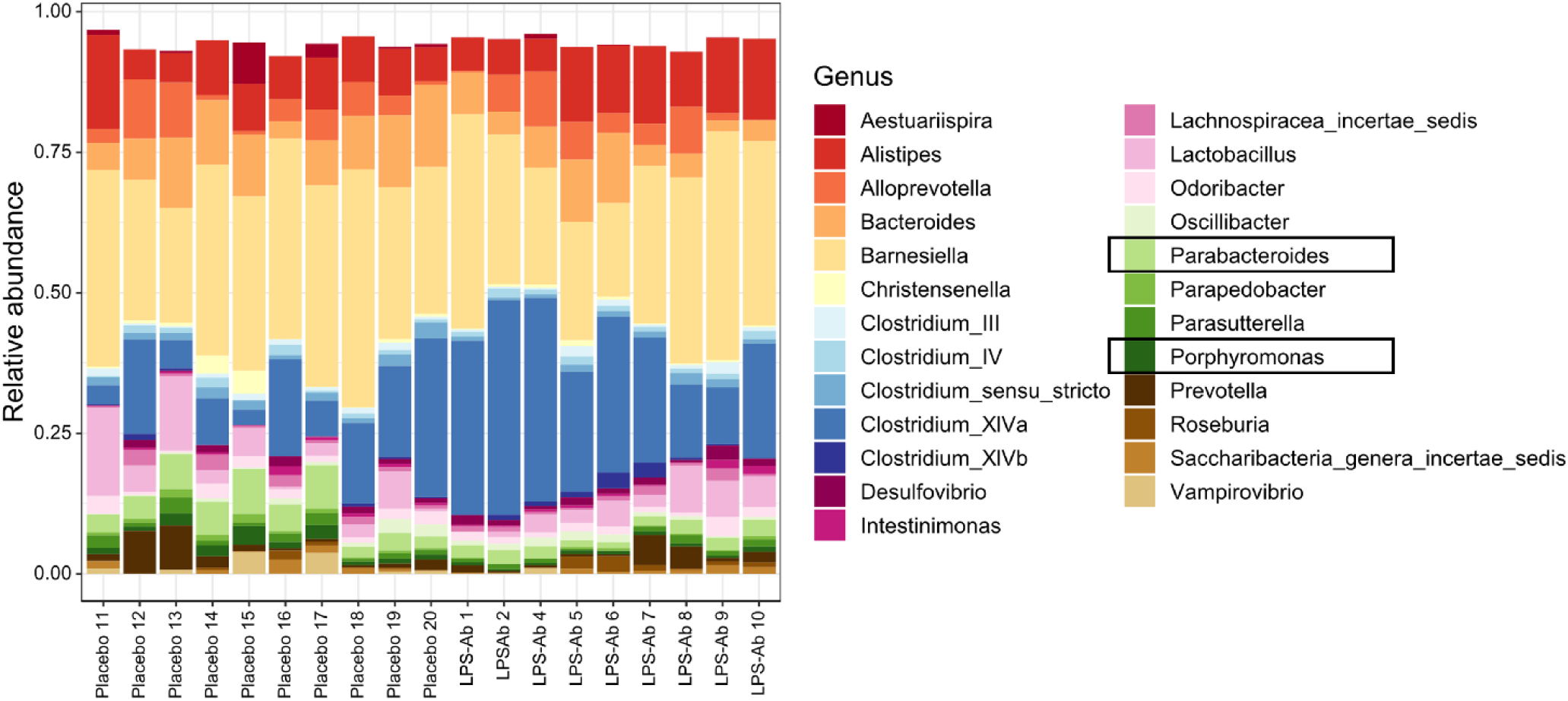
Descriptive bar charts of cecal communities from Travelan® LPS antibody-treated and control communities. OTUs have been colored by genus. Y axis represents % abundance. 78 bacterial genera each with a mean abundance < 0.005% have been excluded for clarity. (Decreased abundance of *Parabacteroides* and *Porphyromonas* visualized here). Although not statistically significant, there was a trend towards an increase in the relative abundance of *Clostridium XIVa* in Travelan® LPS antibody-treated mice.

There was no significant difference in the relative abundances of the OTUs in the colon mucosal samples of the control and mice treated with Travelan® LPS antibody treatment.

## 5 Discussion

The human gut is a major reservoir of lipopolysaccharide (LPS) resulting from shedding of the outer layer of gram-negative bacteria. Elevated LPS levels promote intestinal inflammation through the release of pro-inflammatory cytokines (Rhee, 2014) in conditions such as necrotizing enterocolitis (Leaphart et al., 2007), inflammatory bowel disease (Magro et al., 2017), obesity (Creely et al., 2007) and non-alcoholic fatty liver disease (Henao-Mejia et al., 2012). The immunological activity of IgGs in hyperimmune colostrum raised against a bacterial LPS extract (Imm124-E, Travelan®, Immuron Ltd. Melbourne, Australia) is of increasing relevance to human health due to its ability to directly target specific pathogens, reduce inflammation by preventing translocation of bacterial antigens across the epithelium and inhibit the induction of proinflammatory cytokines (as reviewed by Ulfman et al., 2018).

Here we show that LPS antibodies can engineer fecal and potentially, cecal bacterial communities, without dramatically disrupting the balance and underlying ecology of these GI communities. The effects of colostrum antibodies targeting LPS on the gut microbiome are poorly studied. Therefore, we aimed to assess the onset and magnitude of changes in the richness and diversity of gut microbial populations in mice administered antibodies targeting LPS. Overall, the treatment had minimal impact upon the total number of bacterial cells, bacterial species, community diversity and evenness in a healthy preclinical model.

We first aimed to investigate for changes in GI anatomical parameters following completion of feeding mice milk powder solution (placebo), or a solution containing Travelan® LPS antibody treatment. We demonstrate that administration of antibodies did not affect the length of small intestine or cecal weight in mice. Interestingly, mice fed LPS antibodies showed a strong trend towards longer colons compared to control mice. These findings suggest no adverse change in gastrointestinal anatomy due to dietary supplementation with Travelan® LPS antibodies.

We showed that although bacterial cell number, bacterial species richness or species abundance distribution (evenness) in fecal and mucosal samples remained unchanged following Travelan® LPS antibody colostrum supplementation, bacterial evenness in cecal samples was reduced compared to samples from placebo-treated mice. In addition, treatment with colostrum containing Travelan® LPS antibodies caused a small but significant shift in microbial community structure in fecal, mucosal and cecal samples indicating that microbial communities were altered along the GI tract, which may contribute to subtly altered gastrointestinal function.

To understand which microbes were potentially driving the observed changes in community structure we investigated changes at the level of bacterial OTUs. Using this approach, we revealed shifts in numerous low-abundance bacterial genera, however, many of these are poorly characterized in the GI tract. Minor changes to rare and low abundance species may have a larger impact than anticipated if unique functions provided by these species are disrupted, especially if these species are important members of the ecological core (Risely, 2020). Further exploration of the roles provided by rare species in the GI tract, as well as the relationship between these species is required.

The largest change to the fecal bacterial community in response to Travelan® LPS antibody treatment was an increase in abundance of the genus *Akkermansia. Akkermansia* is involved in maintaining intestinal barrier integrity, reducing inflammation, regulating host metabolism and plays a role in immune tolerance of commensal organisms (Cani & de Vos, 2017; Chelakkot et al., 2018; Derrien et al., 2011; Geerlings et al., 2018). It is widely associated with healthy mucus-associated microbial communities and is a prolific re-colonizer of the GI tract after disturbances such as infection (Geerlings et al., 2018). Based on the findings of the current study, we speculate that Travelan® LPS antibody treatment creates ecological niches for *Akkermansia* to proliferate into by knocking down/removing other bacterial species whose abundances were below the detection threshold of our current statistical approaches.

In Travelan® LPS antibody-treated colonic mucosal communities, an increase of Desulfovibrionales was observed at higher taxonomic levels (i.e., Family and Order). Elevated levels of *Desulfovibrio*, a sulfate-reducing bacteria, are commonly associated with a penetrable mucus phenotype in animals exhibiting gut inflammation (Jakobsson et al., 2015; Rodriguez-Pineiro & Johansson, 2015). While increase in Desulfovibrionales may indicate the presence an inflammatory response (Devkota et al., 2012; Verma et al., 2010) the change may also be attributable to the important metabolic role *Desulfovibrio* plays in the gastrointestinal tract. The utilization of excess hydrogen through the processes of sulfate reduction, methanogenesis and acetogenesis by hydrogen cross-feeders assists in the maintenance of gut homeostasis (Oliphant & Allen-Vercoe, 2019). *Desulfovibrio* is the most dominant group of sulfate-reducing bacteria in the colon (Gibson et al., 1990; Nava et al., 2012); utilizing hydrogen to convert sulfate to sulfide compounds while oxidizing lactate to acetate (Marquet et al., 2009), a beneficial short-chain fatty acid. Interestingly, hydrogen and the products of hydrogen-feeders are associated with both health promoting effects such as mucus layer integrity (Motta et al., 2015; Tomasova et al., 2016), and detrimental health outcomes observed in Parkinson’s disease and inflammatory bowel disease (Chassard et al., 2012; King et al., 1998; Ostojic, 2018; Scheperjans et al., 2015). This apparent contradiction may reflect differences in the microbial species composition involved in hydrogen cycling within individuals. In addition, the sulfate utilized by sulfate-reducing bacteria like *Desulfovibrio* may be derived from dietary sources of sulfur or released from the breakdown of endogenous mucins by other microorganisms, such as *Akkermansia* (Kosciow & Deppenmeier, 2020; Van Herreweghen et al., 2018). Therefore, an increase in Desulfovibrionales without evidence of pathology may merely indicate an increase in substrate availability due to an increased abundance of mucin degraders like *Akkermansia*.

Although several changes were observed in cecal communities due to administration of Travelan® LPS colostrum, only community members of relatively rare abundance were impacted, and the abundance changes were minor. We observed a reduction in cecal microbial community evenness indicating that Travelan® LPS antibody-treated cecal communities have more dominant OTUs. This may be due to the trend for an increase in *Clostridium XIVa* abundance in Travelan® LPS antibody-treated cecal samples, which was marginally, but not statistically significant. These community diversity metrics incorporate both OTU richness and evenness. Our finding that the cecal community diversity was unaltered despite a shift in evenness, suggests that community evenness changes induced by treatment with Travelan® LPS antibodies are minimal, although statistically significant.

Collectively, these data suggest that LPS colostrum treatment administered in the absence of infection does not adversely affect the GI microbial community structure. The ability to target specific bacterial species, without large off-target effects, makes this a promising tool in selective modification of GI microbial composition in diseases with gastrointestinal co-morbidities. Further investigation is needed to determine whether these subtle shifts in abundance translate to a change in the functional attributes of these microbial communities.

## 6 Conclusions

This is the first study to assess the impact of Travelan® LPS antibodies on anatomical and microbial changes in the wild-type non-immune challenged mouse gastrointestinal tract. There was no difference in any of the anatomical measures obtained between groups, however, we did identify small but significant differences in rare bacterial species of mouse gastrointestinal microbial populations following treatment with Travelan® LPS antibodies. Specifically, increases in the abundance of the gut bacterial genera *Akkermansia* were identified in fecal samples as well as reductions in the abundance of eight relatively rare OTUs in the cecal microbial community. The increase in *Akkermansia* may be a beneficial off-target outcome caused by the reduction in rare bacterial species induced by Travelan® LPS antibody treatment. Our results indicate that administering Travelan® LPS antibody treatment to a healthy preclinical model can modulate fecal and potentially cecal bacterial communities without dramatically disrupting the balance or underlying ecology of gastrointestinal communities. Further studies are required to examine the potential of Travelan® LPS antibody treatment to engineer the microbiome to alleviate gut dysbiosis in preclinical models and humans.

## 8 Author Contributions

Conceptualization, EH-Y, AF and GB; methodology, GB, MM, EH-Y, RG; formal analysis, MM, RG, JW; investigation, MM, RG; bioinformatics, JW and RG; EH-Y, JW, AF, RG, GB and MM wrote the manuscript, funding acquisition, EH-Y and AF. All authors have read and agreed to the published version of the manuscript.

## 10 Data Availability Statement

The datasets presented in this study can be found in online repositories. The names of the repositories are as follows: NCBI BioProject (accession number PRJNA785752), RMIT University figshare repository (DOI: 10.25439/rmt.17141858).

