## Supplementary Material for "Hyperimmune bovine colostrum containing lipopolysaccharide antibodies (Imm124-E) has a non-detrimental effect on gut microbial communities in unchallenged mice"

Supplementary Table 1. Gastrointestinal tract anatomical measures normalized to individual body weight.

| Measurement (normalized to body weight) | Placebo  (mean ± SEM) | LPS-Ab treatment (mean ± SEM) | n  (per group) | t-score | p value |
| --- | --- | --- | --- | --- | --- |
| Cecum weight | 0.025 ± 0.001 | 0.028 ± 0.002 | 10 | *t*_(18)_= 1.544 | 0.140 |
| Small intestine length | 2.02 ± 0.037 | 1.94 ± 0.104 | 10 | *t*_(18)_= 0.7689 | 0.452 |
| Colon length | 0.339 ± 0.010 | 0.379 ± 0.017 | 10 | *t*_(18)_= 2.041 | 0.056 |

Values are the ratio mean ± SEM of the number of mice indicated. P values are for LPS Ab-treated mice compared to placebo-treated mice.

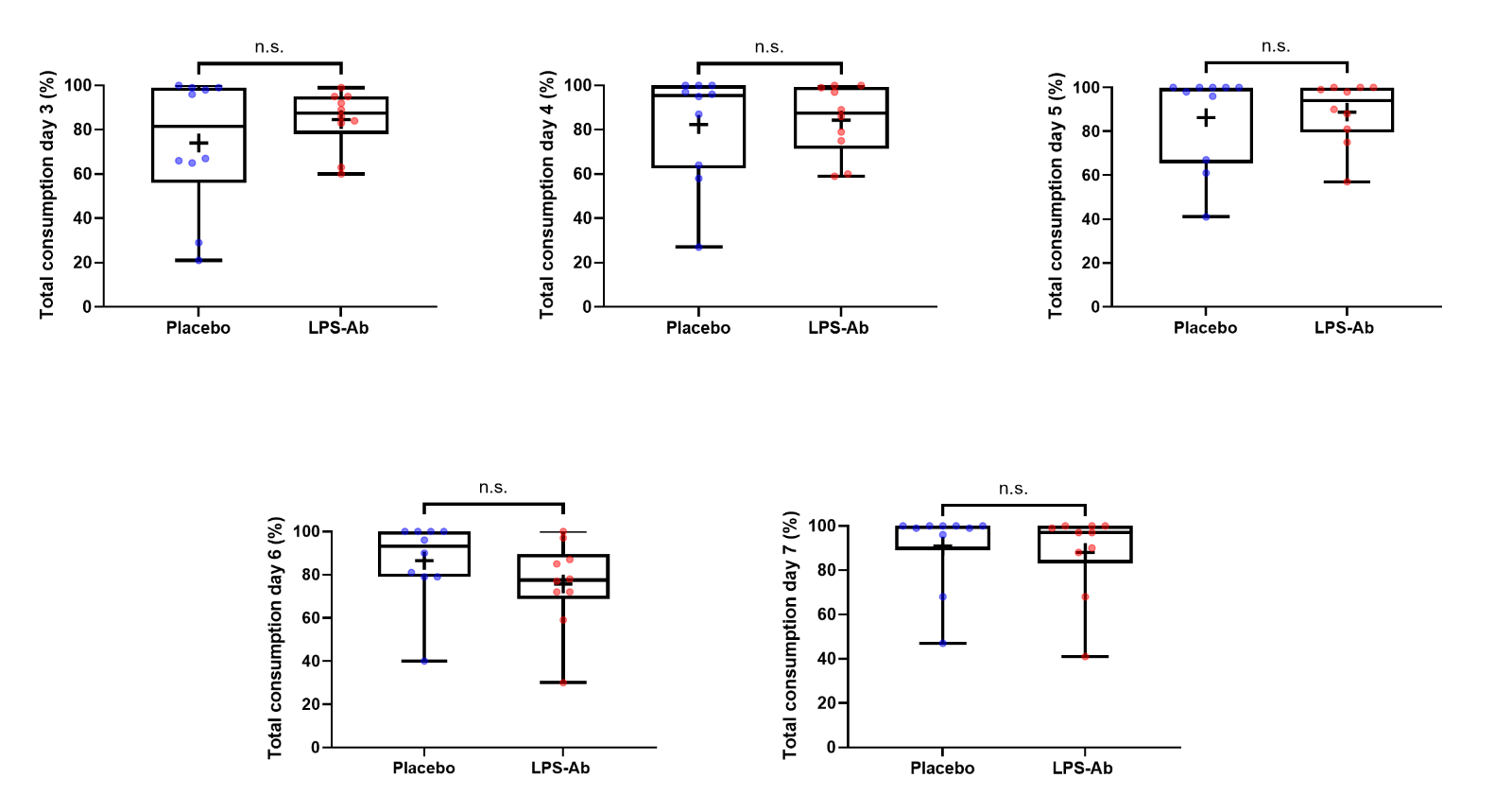

**Supplementary Figure 1. Percentage consumption of placebo/ Travelan® LPS antibody treatment (LPS-Ab) (total volume = 4 ml) per mouse per day. Day 3 represents the first day of feeding the mice with the milk powder placebo solution or Travelan® LPS antibody treatment (n=10 placebo, n=10 LPS-Ab), P>0.05 for all comparisons. Groups were compared using a two-tailed Student’s *t*-test (GraphPad Prism v9.0.1). Placebo: Promilk 85, Tatura Milk Industries Ltd, Australia); LPS-Ab: Travelan® LPS antibody treatment****.** **Boxplots represent median, interquartile range and range of the data; cross represents the mean.**

Supplementary Table 2. Percentage consumption of offered 4 mL of placebo or Travelan® LPS antibody treatment per mouse.

| Solution consumption  (% of total) | Placebo  (mean ± SEM) | LPS-Ab treatment (mean ± SEM) | n  (per group) | t-score | p value |
| --- | --- | --- | --- | --- | --- |
| Day 3 | 74.0 ± 9.44 | 84.6 ± 4.18 | 10 | *t*_(18)_= 1.027 | 0.318 |
| Day 4 | 82.4 ± 7.83 | 84.4 ± 4.99 | 10 | *t*_(18)_= 0.2155 | 0.832 |
| Day 5 | 86.3 ± 6.86 | 88.8 ± 4.51 | 10 | *t*_(18)_= 0.3046 | 0.764 |
| Day 6 | 86.5 ± 5.90 | 75.7 ± 6.38 | 10 | *t*_(18)_= 1.243 | 0.230 |
| Day 7 | 90.9 ± 5.80 | 88.0 ± 6.08 | 10 | *t*_(18)_= 0.3452 | 0.734 |

Values (%) are the mean ± SEM of the number of mice indicated. P values are for LPS Ab-treated mice compared to placebo-treated mice.

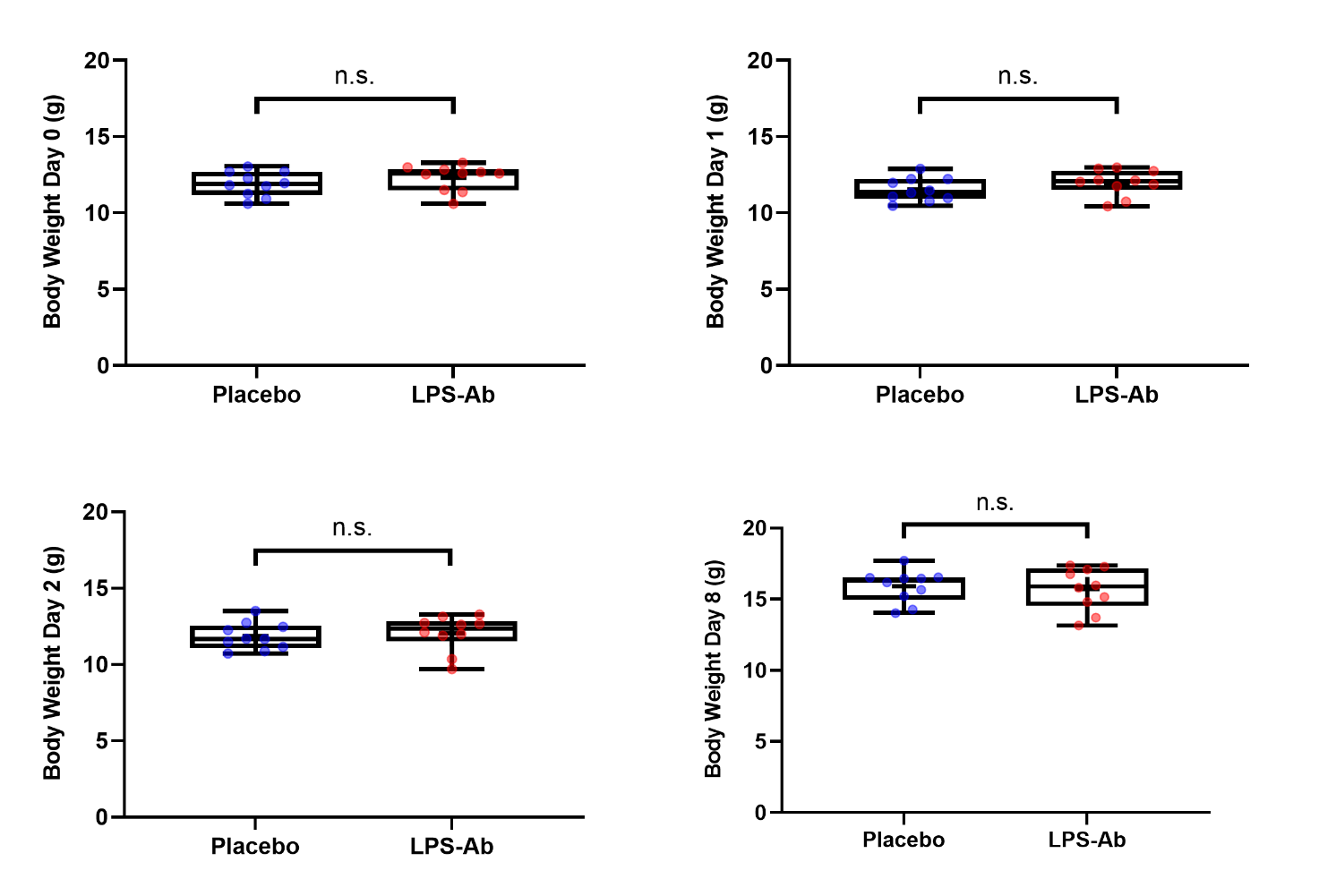

**Supplementary Figure 2. All mice were weighed daily for 3 consecutive days before feeding and weighed once after feeding with placebo/ Travelan® LPS antibody treatment (LPS-Ab). Day 0 represents one day before the pre-feeding pellet collection, day 1 and day 2 represent the first and second day of pre-feeding pellet collection, respectively and day 8 represents the day of the tissue collection (n=10 placebo, n=10 LPS-Ab). P>0.05 for all comparisons. Groups were compared using a two-tailed Student’s *t*-test (GraphPad Prism v9.0.1). Placebo: Promilk 85, Tatura Milk Industries Ltd, Australia); LPS-Ab: Travelan® LPS antibody treatment. Boxplots represent median, interquartile range and range of the data; cross represents the mean.**

Supplementary Table 3. Body weights of mice before the commencement and on termination of the feeding protocol.

| Body weight  (g) | Placebo  (mean ± SEM) | LPS-Ab treatment (mean ± SEM) | n  (per group) | t-score | p value |
| --- | --- | --- | --- | --- | --- |
| Day 0 | 11.9 ± 0.255 | 12.3 ± 0.269 | 10 | *t*_(18)_= 1.066 | 0.301 |
| Day 1 | 11.5 ± 0.241 | 12.0 ± 0.267 | 10 | *t*_(18)_= 1.206 | 0.244 |
| Day 2 | 11.9 ± 0.278 | 12.0 ± 0.369 | 10 | *t*_(18)_= 0.4004 | 0.694 |
| Day 8 | 15.9 ± 0.357 | 15.7 ± 0.472 | 10 | *t*_(18)_= 0.3227 | 0.751 |

Values (g) are the mean ± SEM of the number of mice indicated. P values are for LPS Ab-treated mice compared to placebo-treated mice.

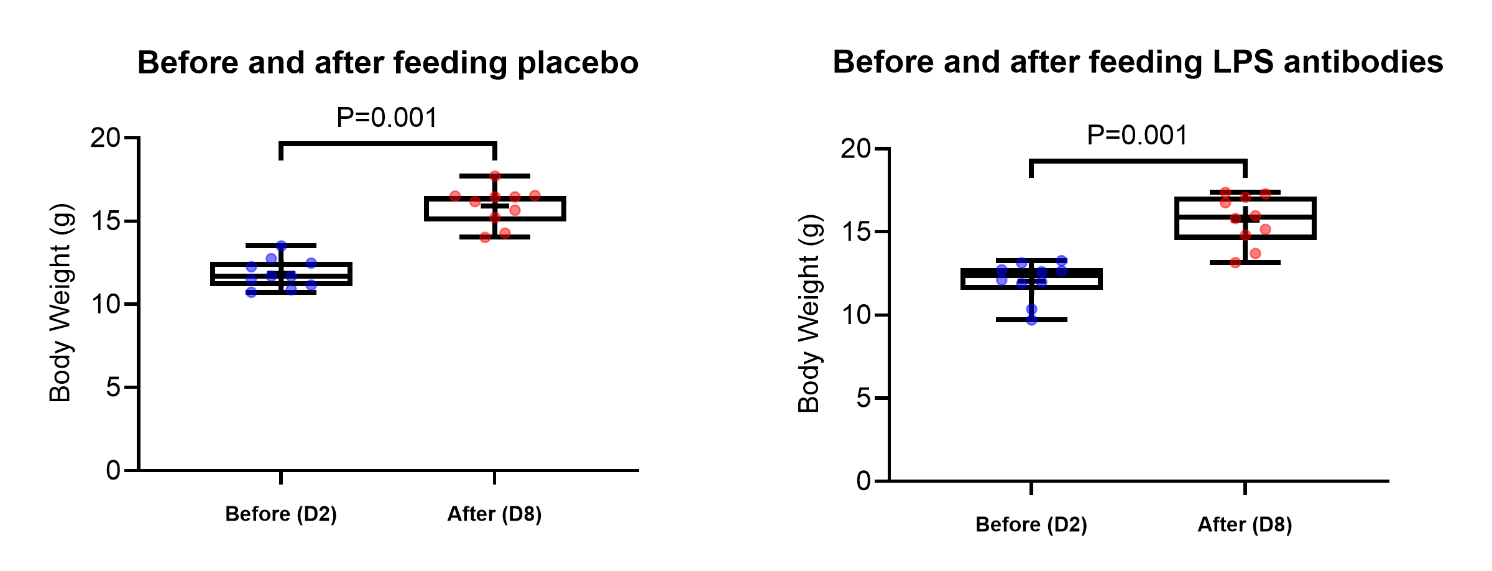

**Supplementary Figure 3. Body weights of mice before and after feeding with placebo/** **Travelan® LPS antibody treatment (LPS-Ab). Day 2 represents the day before the feeding protocol was initiated and day 8 represents the day of the tissue collection (n=10 placebo, n=10 LPS-Ab). The results indicate a significant difference in body weight (g) before commencement of the feeding protocol (****11.86 ± 0.278 g, n=10) and at the end of the experiment (15.90 ± 0.357 g, n=10); [t(9) = 20.47, p = .001] for mice fed the placebo solution. A significant difference in body weight (g) was also observed for the Travelan® LPS antibody treatment group before commencement of the feeding protocol (****12.04 ± 0.369 g, n=10) and at the end of the experiment (15.71 ± 0.472 g, n=10)****; [t(9) = 7.976, p = .001].. Groups were compared using a two-tailed paired *t*-test (GraphPad Prism v9.0.1). Placebo: Promilk 85, Tatura Milk Industries Ltd, Australia); LPS-Ab: Travelan® LPS antibody treatment. Data are presented as mean ± SEM. Boxplots represent median, interquartile range and range of the data; cross represents the mean.**

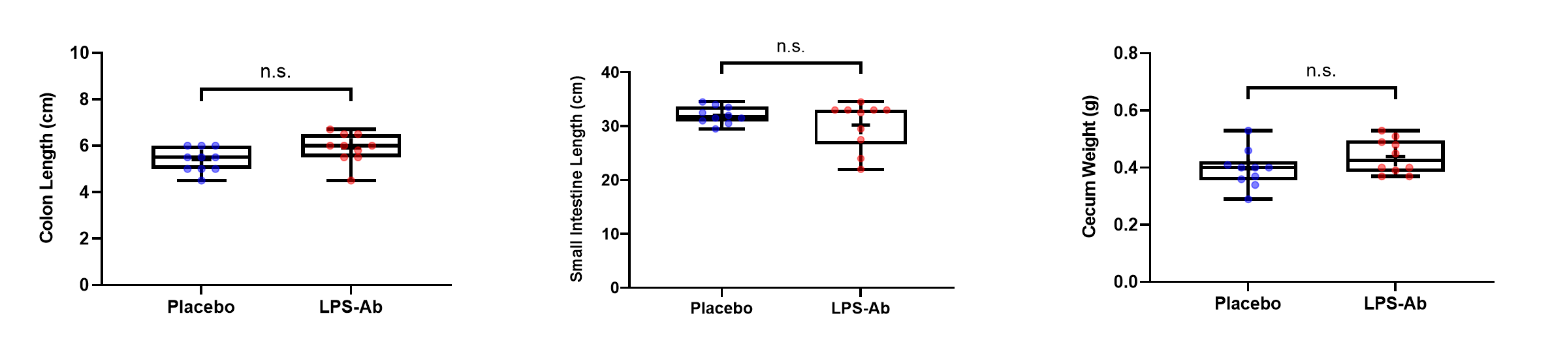

**Supplementary Figure 4.** **Gastrointestinal measurements not normalized against body weight for placebo and Travelan® LPS antibody treatment (LPS-Ab) mice (n=10 placebo, n=10 LPS-Ab). P>0.05 for all comparisons, although these was a trend towards an increase in colon length for Travelan® LPS antibody treatment mice (P=0.070). Groups were compared using a two-tailed Student’s *t*-test (GraphPad Prism v9.0.1). Placebo: Promilk 85, Tatura Milk Industries Ltd, Australia); LPS-Ab: Travelan® LPS antibody treatment. Boxplots represent median, interquartile range and range of the data; cross represents the mean.**

Supplementary Table 4. Gastrointestinal measurements not normalized against body weight.

| Measurement | Placebo  (mean ± SEM) | LPS-Ab treatment (mean ± SEM) | n  (per group) | t-score | p value |
| --- | --- | --- | --- | --- | --- |
| Colon length (cm) | 5.40 ± 0.52 | 5.90 ± 0.64 | 10 | *t*_(18)_= 1.924 | 0.070 |
| Small intestine length (cm) | 32.1 ± 1.59 | 30.2 ± 4.33 | 10 | *t*_(18)_= 1.269 | 0.221 |
| Cecum weight (g) | 0.396 ± 0.07 | 0.439 ± 0.06 | 10 | *t*_(18)_= 1.527 | 0.144 |

Values are the mean ± SEM of the number of mice indicated. P values are for LPS Ab-treated mice compared to placebo-treated mice.
